# Effects of Superhydrophobic Sand Mulching on Evapotranspiration and Phenotypic Responses in Tomatoes (*Solanum lycopersicum*) under Normal and Reduced Irrigation

**DOI:** 10.1101/2021.10.26.465960

**Authors:** Kennedy Odokonyero, Adair Gallo, Vinicius Dos Santos, Himanshu Mishra

## Abstract

Irrigated agriculture in arid and semi-arid regions is a vital contributor to the global food supply; however, these regions endure massive evaporative losses that are compensated by unsustainable freshwater withdrawals. Plastic mulches have been used to curtail evaporation, improve water-use efficiency, and ensure food–water security, but they are non-biodegradable and their disposal is unsustainable. We recently developed superhydrophobic sand (SHS), which comprises sand grains with a nanoscale wax coating that could offer a more sustainable mulching solution. Here, the effects of adding a 1.0 cm-thick layer of SHS mulch on the evapotranspiration and phenotypic responses of tomato (*Solanum lycopersicum*) plants are studied under normal and reduced irrigation. Under both irrigation regimes, SHS mulching suppressed evaporation and enhanced transpiration by 78% and 17%, respectively relative to the bare soil. Overall, SHS mulching enhanced root xylem vessel diameter, stomatal aperture, stomatal conductance, and chlorophyll content index by 21%, 25%, 28%, and 23%, respectively. Total fruit yields, total dry mass, and harvest index increased in SHS-mulched plants by 33%, 20%, and 16%, respectively than in bare soil. These findings demonstrate the potential of SHS to boost irrigation efficiency in water-limited environments and provide mechanistic insights behind yield enhancement by SHS mulching.

## INTRODUCTION

Irrigated agricultural lands are of vital importance, as they represent 20% of the total cultivated land but contribute over 40% of the global food production (WWAP, 2009). Irrigated agriculture has been expanding to feed the growing global population; for example, India, China, and Brazil have witnessed a 30% (Jain et al., 2020), 52% (Zhu et al., 2013), and 52% (Carvalho et al., 2020) increase in cultivated areas under irrigation in the last 50 years, respectively. Further, between 2012 and 2030, the global irrigated land area is expected to increase by over ≈30%, from 310 Mha to 402 Mha (Darko et al., 2016). These food production trends are sustainable only when freshwater is abundant. However, groundwater resources have been extensively exploited for irrigation in densely populated arid and semi-arid regions and are thus rapidly declining (Steward et al., 2013, Watto et al., 2018b, Watto et al., 2018a, Scanlon et al., 2012, Famiglietti, 2014). Therefore, water consumption patterns in irrigated agriculture must be studied, and new technologies must be created to achieve global water–food security.

Water applied to topsoil is lost via evaporation, transpiration, and percolation (Sutanto, 2012). High temperatures and dry winds in arid and semi-arid regions lead to substantial evaporative and transpiration losses (Al-Naizy, 2012, Balugani et al., 2017), whereas percolation is due to the poor water-holding capacity of sandy soils (Lehmann et al., 2019, Or and Lehmann, 2019). These water losses are compensated via irrigation that, due to the sheer size of agricultural operations, claims the lion’s share (approximately ≈80%) of global annual freshwater consumption (2500 km^3^/yr) (Shiklomanov, 2000, Hoekstra and Mekonnen, 2012). Whereas evaporation and percolation do not directly contribute to photosynthesis during plant growth, transpiration is crucial in maintaining the optimal temperature required for plants’ metabolic processes (Grill and Ziegler, 1998, Hetherington and Woodward, 2003). Transpiration thus cannot be reduced while high crop productivity is maintained without genetic engineering. Percolation could be reduced by applying a water barrier layer beneath the plant root zone, but this process is labor-intensive and expensive (Nkurunziza et al., 2019). Thus, many researchers have focused on reducing evaporative losses.

Together, evaporation and transpiration constitute evapotranspiration (ET), a process that plays an important role in irrigation and agriculture practices (Hatfield and Dold, 2019, Rawitz and Hadas, 1994). The importance of transpiration is reflected in the water-use efficiency (WUE) of plants, i.e., the ratio of the total biomass (root and shoot) formed to the cumulative water transpired (Kadam et al., 2015), and the transpiration efficiency (TE), i.e., the ratio of the shoot biomass produced to the cumulative water transpired (Vadez et al., 2014). The TE represents an aspect of WUE that depends on the water-conducting (i.e., hydraulic) potential of a plant to facilitate shoots’ physiological processes that enhance biomass under different soil water scenarios(Vadez et al., 2014). Therefore, the TE is of interest in water-scarce environments where crops are most sensitive to soil moisture (Condon et al., 2002, Sinclair, 2012). Transpiration also influences the plants’ reproductive efficiency, defined by the harvest index (HI) entailing the ratio of total yield to total vegetative biomass produced, pinpointing the plant resources invested in yield production (Porker et al., 2020, Unkovich et al., 2010). Hence, whereas water uptake and efficient use of water are critical for plant growth and biomass accumulation, the development and growth of reproductive organs determines the HI, which is critical to crop yield (Hammer et al., 2021).

Notably, evaporation is a wasteful component of ET and should be minimized. Mulching, i.e., applying a vapor diffusion barrier on the topsoil to curtail water evaporation, has been proven to enhance soil moisture content and provide enhanced transpiration (Moitra, 1996, Zhang et al., 2018, Farzi et al., 2017), plant biomass, and yields (Ramalan, 2000, Zhang, 2017a, Mukherjee, 2010). Mulching has been demonstrated to enhance leaf chlorophyll content (Wang et al., 2015), photosynthesis (Niu et al., 2020, Zhang et al., 2019), and TE (Balwinder-Singh et al., 2011), as well as improve plant root growth and roots’ architectural and anatomical properties, such as root diameter and late xylem vessel diameter, which improves water and nutrient uptake from the soil (Zhan et al., 2019, Larsson and Jensen, 1996). Low-density polyethylene sheets approximately 0.1 mm thick have been used extensively for mulching in developed countries (Kasirajan and Ngouajio, 2012). Their application in the developing world is also on the rise, e.g., consumption in China is set to exceed two million tons per year by 2024 (Liu et al., 2014, Wang et al., 2013). Despite their benefits (Zhang, 2017b, Zhang et al., 2019, Balwinder-Singh et al., 2011, Yin W, 2019, Li, 2004), plastics are non-biodegradable, and their eventual disposal into landfills is unsustainable (Kasirajan and Ngouajio, 2012, Halley et al., 2001). Recent findings on the leaching of phthalates from plastic mulches into soils and their adverse effects on the soil quality and microbial activity over the long term are also concerning (Shah and Wu, 2020). Therefore, sustainable mulching technologies are needed.

We recently developed superhydrophobic sand (SHS) mulch technology (Mishra et al., 2017, Gallo Jr et al., 2021a), a nature-inspired material comprising common sand gains or sandy soils coated with a nanoscale layer of paraffin wax (wax to sand mass ratio is 1:1000). The micro- and nanoscale surface roughness of the sand grains/particles and the hydrophobic nature of wax induce superhydrophobicity (Arunachalam et al., 2019). When laid on topsoil with a sub-surface irrigation system, a 5–10 mm-thick SHS layer can insulate the wet soil from solar radiation and dry air, thereby acting as a diffusion barrier, which reduces evaporation (Gallo Jr et al., 2021b). Multi-year field trials of SHS on tomatoes (*Solanum lycopersicum*), wheat (*Triticum aestivum*), and barley (*Hordeum vulgare*) under naturally arid land conditions in Saudi Arabia have revealed significant enhancements in plant growth and yields (Gallo Jr et al., 2021a). Compared with plastic mulch, SHS is cheaper and more environmentally friendly, as it is made from readily abundant sands and paraffin wax that is easily degraded by soil microbes (Marino, 1998, Roper, 2004). After nine months of use, the wax is decomposed by microbial activity and solar radiation and incorporated into the soil after the crop cycle without affecting soil microbial compositions, obviating landfilling (Gallo Jr et al., 2021a). Despite these promising field results, quantitative insights into the effects of SHS on ET and the phenotypic traits responsible for yield enhancement are lacking.

This work therefore aims to quantify the effects of mulching with SHS on tomato plants by pinpointing the ET dynamics and consequent plant phenotypes, including plant height, stomatal conductance, stomatal pore size (aperture), leaf chlorophyll content, fruit yields, HI, TE, fresh mass, dry mass, xylem vessel and root diameter. These effects are considered in plants grown in controlled growth chambers under normal (**N**) and reduced (**R**) irrigation scenarios at 100% field capacity and 50% of field capacity, respectively.

## MATERIALS AND METHODS

### Plants and SHS mulch

The tomato plants (*Solanum lycopersicum*), Seminis variety (St. Louis, Missouri, US), were purchased from local seed stores in Jeddah, Saudi Arabia. The SHS was produced from common sand and paraffin wax following the optimized protocol detailed in previous work (Mishra et al., 2017, Gallo Jr et al., 2021a); readers are referred to these works for details on SHS production. Briefly, sand was added to an organic solvent containing dissolved wax. The solvent was then removed by changing the pressure and temperature and condensed for reuse, leaving behind the SHS.

### Plant growth conditions, treatment, and experimental design

The tomato seeds were sown in plastic trays using a potting mix from Stender AG (Schermbeck, Germany) and grown in Percival growth chambers. After four weeks, the seedlings were transplanted into pots 1,870 cm^3^ in volume (15 cm top diameter × 10.5 cm bottom diameter × 14.5 cm height) containing approximately 2.4 kg of local sandy soil. The pots were watered via sub-surface irrigation to two field capacity levels every two days: 100% field capacity for **N** irrigation (i.e., the maximum soil moisture content after drainage of excess water from fully saturated potted soil) and 50% of this field capacity for **R** irrigation. The irrigation level of each pot was determined gravimetrically. Thirty-two pots were prepared, including 16 with plants and 16 without plants, the latter of which were used to quantify evaporative losses.

The pots were separated into two groups: those comprising SHS and those containing only bare soil. In each group, half of the pots were subjected to **N** irrigation while the other half was maintained under **R** irrigation. For pots containing SHS, a 1.0 cm-thick layer of SHS was applied to the potting soil (approximately 265 g SHS/pot). Thus, there were four treatment combinations (SHS-**N**, bare soil-**N**, SHS-**R**, and bare soil-**R)**, each with two replicates with and without plants. All pots were subjected to complete randomization in a 2 × 2 factorial design involving two experimental factors (i.e., soil mulching and irrigation regime), each with two levels and four replicates. During the growth period, nutrient solutions were applied to each pot every two weeks comprising in rates per kg of dry soil: 200 mg N/kg, 250 mg P/kg, 200 mg K/kg, 150 mg Ca/kg, 30 mg S/kg, 2 mg Cu/kg, 4 mg Zn/kg, 3 mg Mn/kg, 0.5 mg B/kg, and 0.25 mg Mo/kg. The plants were grown for 98 days under a 14/10 h light/dark photoperiod using artificial fluorescent lighting at 800 μmol m^−2^ s^−1^ of photosynthetically active radiation, a day/night temperature of 28/20°C ± 2°C, and 60 ± 2% air relative humidity.

### Evapotranspiration (ET)

The ET was partitioned into evaporation and transpiration by performing gravimetric measurements of pots every two days until the final harvest. The water loss from each pot was monitored, and water was added to compensate for the water lost. The daily ET was recorded as the total water lost from each pot containing plants between the time of irrigation to the time of weighing as:

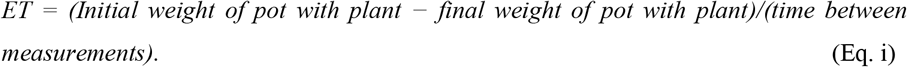

### Evaporation

Under the assumption that the pots with and without plants experienced a similar amount of evaporation, the daily water loss by evaporation was estimated using pots without plants as:

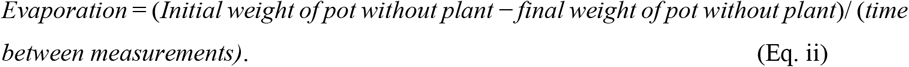

### Transpiration

The daily transpiration was determined as:

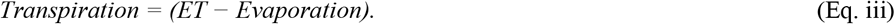

### Soil water content (SWC)

The total available soil water content (SWC; i.e., the field capacity) was maintained at a relatively constant level by compensating for the water lost through ET after each gravimetric measurement scheduled at two-day interval, as has been done by prior researchers (Ray and Sinclair, 1998, Pellegrino et al., 2004).

### Stomatal conductance and leaf chlorophyll content

The leaf stomatal conductance was determined using an AP4 Porometer (Delta T, Cambridge, UK). Measurements were performed on three young but fully expanded leaves once a week (between 10:00 and 12:00), and the mean conductance for each treatment combination was calculated. The leaf chlorophyll content index (CCI) was measured using a CCM-200 Chlorophyll Content Meter (Optic-Sciences, Inc. Hudson NH03051, USA), with measurements being performed on three young but fully expanded leaves.

### Leaf stomatal pore and root anatomical features

Microscopic analyses of leaf stomata and root anatomic structures were performed on samples collected on the final date of harvest. Two fully expanded upper leaves were cut from each plant (between 11:00 and 11:30) and immediately immersed in 70% ethanol inside 50 ml centrifuge tubes until they were analyzed. After two weeks, the preserved leaf tissues were washed with deionized water three times, added to 40 ml of concentrated sodium hypochlorite, and left to stay for approximately 4 hours. The leaf tissues were re-washed with deionized water, and fresh 70% ethanol was again added onto the samples. This clearing process removed the chlorophyll from the leaves and rendered them white in appearance. The stomata of the cleared leaves were then observed using a digital microscope (Leica DVM6) equipped with the tools to measure the length and width of the stomatal pores. The total stomatal aperture area was calculated using the formula for the area of an ellipse, as

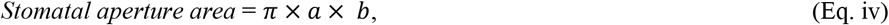

where *a* and *b* represent the semi-major axis (or radius) and semi-minor axis (or width) of the ellipse, respectively.

Root samples 10 cm in length from the growing tip were cut using a razor, washed in deionized water, and fixed in 70% ethanol until their microscopic analysis. Two hand-cut root tissues were made per plant and their cross-sections were analyzed using the same digital microscope to measure diameter of the xylem vessels and the total root.

### Plant growth, fruit yield, and biomass

The plant height was measured every week during the 98–day experimental period. Tomatoes were harvested and weighed at the time of harvest. During the final harvest, the shoots and roots of each plant were weighed, and then put in paper bags and oven-dried at 105 ºC for four days to determine their dry mass.

### Harvest index (HI) and transpiration efficiency (TE)

After harvest, the HI was calculated for each plant as the ratio of total fruit yield (fruit mass per plant) to total fresh mass (shoot and roots) per plant, as:

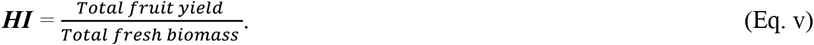

The TE of each plant was calculated as the ratio of dry shoot biomass in grams to the amount of water transpired in kilograms, as:

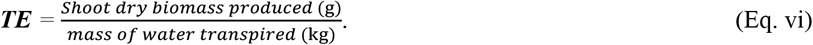

### Data analysis

Data analysis and graphical presentations were performed using Origin Pro Software (2019 version). A three-way analysis of variance (ANOVA) was used to analyze the effects of soil mulching, irrigation regimes, growth stage (i.e., time), and their interactions on ET. For the variables averaged across time (i.e., time-independent), a two-way factorial ANOVA was performed to determine the effects of soil mulching, irrigation regimes, and their interactions. In each analysis, mean comparisons were done using the Tukey test at the *p* < 0.05 level of statistical significance.

## RESULTS

### Plant growth

Representative snapshots of experimental plants grown under each condition (i.e., bare soil or soil containing SHS and **N** or **R** irrigation) at different weeks during their growth, from transplanting (week 0) to fruit ripening (week 12), are shown in **Fig. 1**. Overall, the plants grown in soils containing SHS were larger and appeared healthier than those grown in bare soil.

**Fig. 1.**
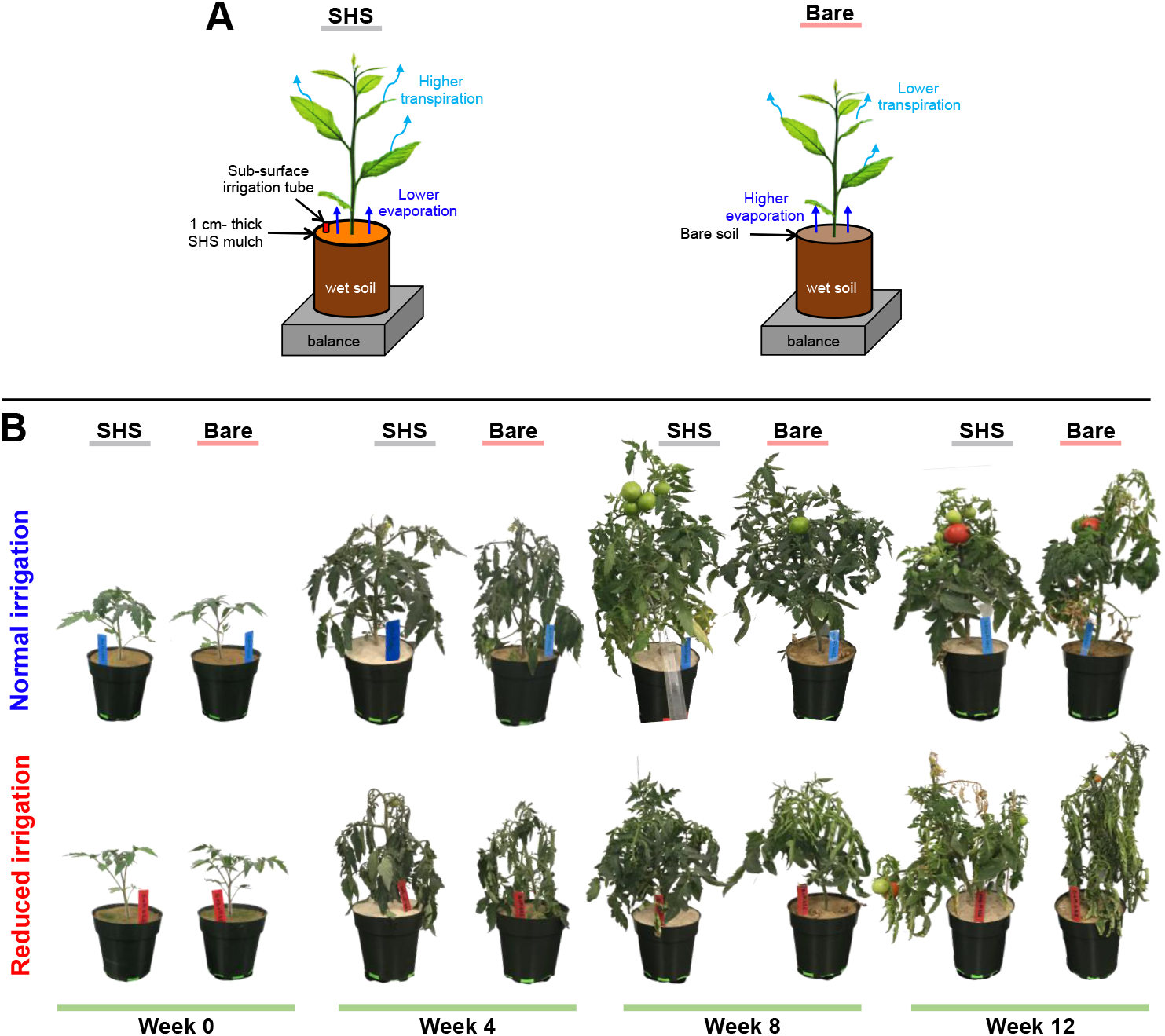
Effects of SHS mulching on plants. (**A**) Schematics of the evaporation and transpiration pathways and their impacts on plants. (**B**) Representative photographs of tomato plants grown in controlled growth chambers mulched with 1-cm-thick superhydrophobic sand (SHS) and bare soil plants under normal (**N**) and reduced (**R**) irrigation. Tomato plants were transplanted and grown for 98 days until the final fruit and biomass harvest. Each treatment combination involved four (*n* = 4) replicate plants.

### Soil water content

The daily SWC corresponding to each irrigation level is detailed in **Fig. 2A**. For both **N** and **R** irrigation strategies (i.e., 100% and 50% field capacity, respectively), the SWC was slightly increased during days 40–98 to compensate for the increase in plant size and water demand. The slight fluctuations in SWC below the initial field capacities observed at five occasions (i.e., between 30–50 days) represent five days when pot weights were recorded without adding water until the following day due to unavoidable circumstances. However, the plants were not severely stressed on these occasions, as the available SWC was enough to meet the ET demand of the plants until the next day when water was added to the pots. When averaged across both mulched and bare soils, the mean SWC was 0.62 ± 0.01 and 0.33 ± 0.01 kg/pot under **N** and **R** irrigation, respectively. This result also demonstrates that plant growth could be maintained under 50% of normal irrigation (i.e., the **R** irrigation strategy).

**Fig. 2.**
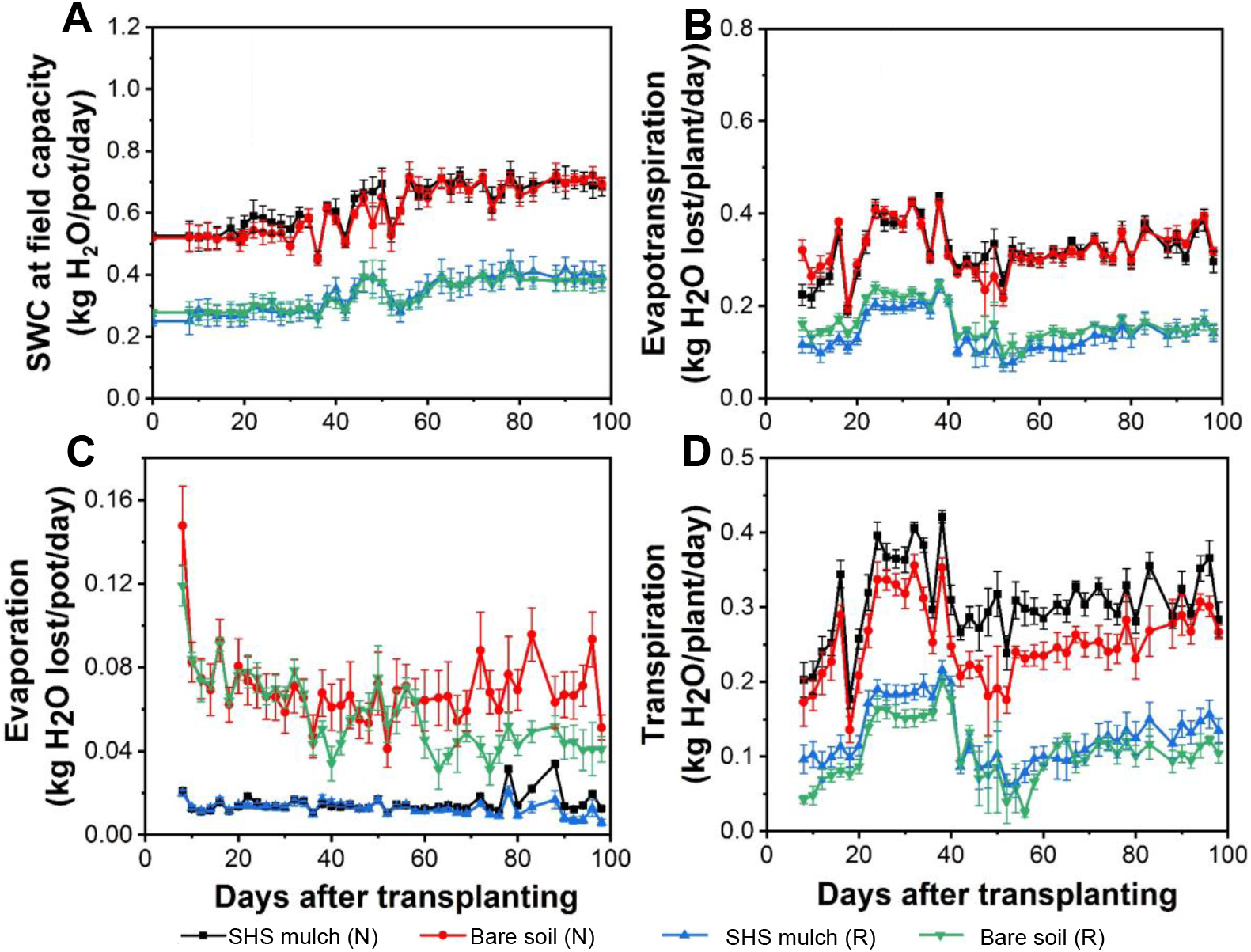
(**A**) Soil water content (SWC) in pots maintained at field capacity for each treatment and irrigation regime, (**B**) daily changes in ET, (**C**) evaporation from the soil, and (**D**) transpiration between SHS-mulched and bare soils under **N** and **R** irrigation strategies throughout plant growth. Each data point is a mean of four (*n* = 4) replicates; error bars are standard errors (±SE) of the means.

### Evapotranspiration

The daily ET significantly differed between the **N** and **R** irrigation scenarios and accounted for approximately 50% of the daily SWC on average, as shown in **Fig. 2B**. Under **R** irrigation, ET differed between mulched and bare soils, notably during days 10–35 and 45– 75; no significant differences were found under **N** irrigation.

### Soil evaporation and transpiration

Throughout the growth period, the daily evaporation was significantly lower from soil containing SHS than bare soil under **N** and **R** irrigation, as shown in **Fig. 2C**. Consequently, plants in mulched soil had higher daily transpiration than in bare soil under both irrigation scenarios, as shown in **Fig. 2D**.

### Cumulative evapotranspiration

There was no significant difference in the cumulative ET loss between mulched and bare soils under **N** irrigation; both had a mean cumulative ET of 13.9 ± 0.7 kg/plant over the 98 days of observation, as shown in **Fig. 3A**. Under **R** irrigation, the cumulative ET was 13% higher in SHS-mulched soil in than bare soils (with mean values of 6.97 ± 0.57 and 6.06 ± 0.78 kg/plant, respectively).

**Fig. 3.**
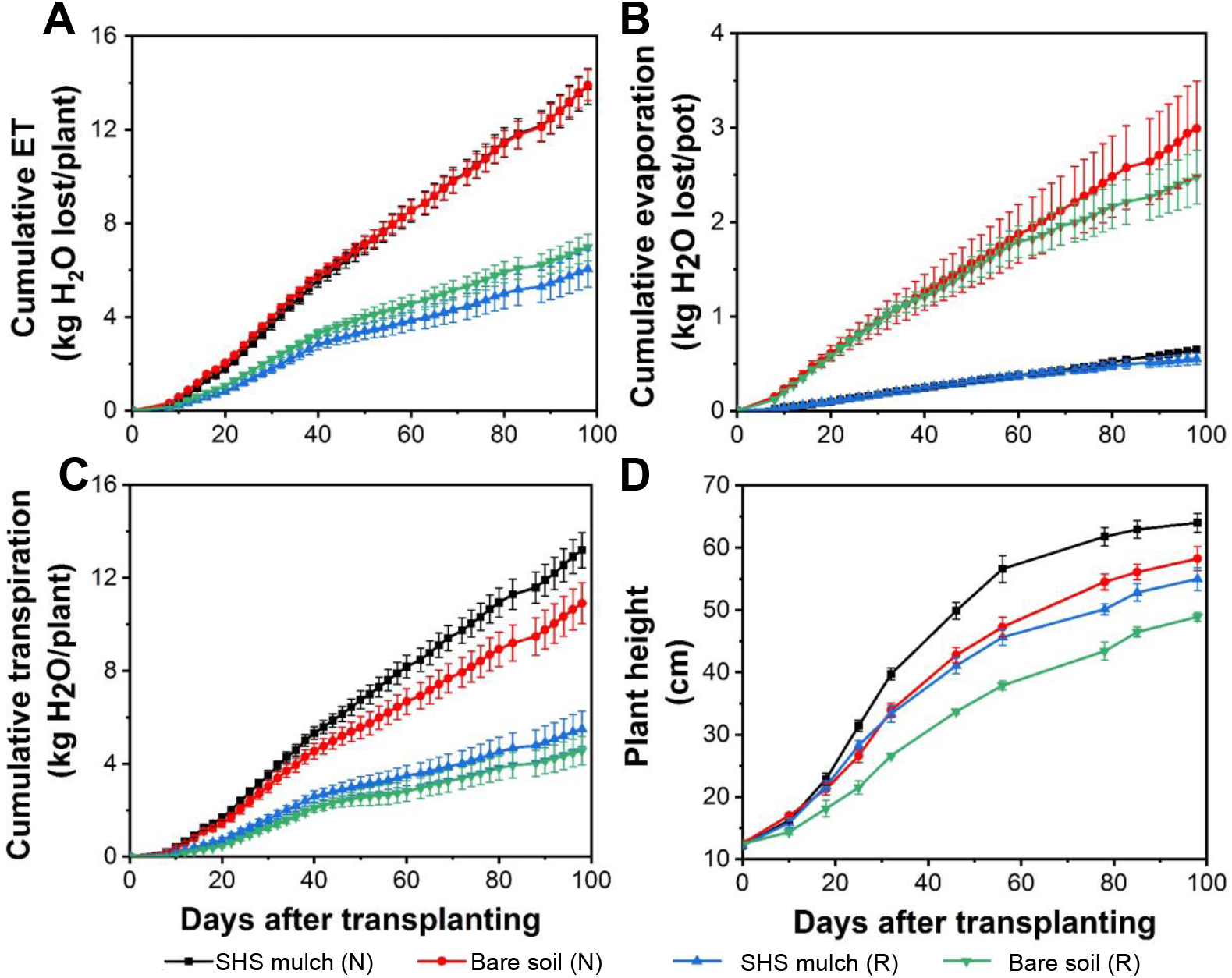
(**A**) Cumulative evapotranspiration (ET), (**B**) evaporation, (**C**) transpiration, and (**D**) plant height during plant growth under contrasting soil mulching treatments (i.e., SHS-mulched vs. bare soil) and irrigation regimes (i.e., **N** and **R**). Each data point is a mean of four (*n* = 4) replicates; error bars represent the ±SE of the mean.

### Cumulative soil evaporation and transpiration

In SHS-mulched soils, that total cumulative evaporation (and transpiration) accounted for 5% (95%) and 9% (91%) of the total ET under **N** and **R** irrigation, respectively; in bare soils, evaporation (and transpiration) accounted for 21.5% (78.5%) and 31.5% (65.5%) of the total ET under **N** and **R** irrigation, respectively (**Figs. 3B and 3C**). Overall, SHS mulching suppressed evaporation and enhanced transpiration by the same magnitude of 78% and 17%, respectively under both **N** and **R** irrigation relative to the bare soils.

### Plant height

Throughout the growth period, the maximum plant height in SHS-mulched soil significantly exceeded that in bare soil by 9% (*p* = 0.005) and 11% (*p* < 0.05) under **N** and **R** irrigation, respectively (**Fig. 3D**). In particular, a more rapid increase in the plant height was observed during 10–45 days after transplanting; after 70 days, plant height started to level off.

### Dependence of evapotranspiration on plant growth stage

The daily mean ET, evaporation, and transpiration during days 0–30, 31–60, and 61–98 are compiled in **Table 1**. Overall, the mean ET, evaporation, and transpiration increased with time; significant differences were present in the mean ET and transpiration between each timeframe (*p* < 0.05). However, no significant difference was found in the mean evaporation from bare soil with time under **N** irrigation (*p* > 0.05), as evidenced by the three-way ANOVA results. Significant two-way interactions were present between mulching and irrigation regimes on transpiration (*p* < 0.05) and between irrigation regime and growth stage on ET, evaporation, and transpiration (*p* < 0.05).

**Table 1.** Effects of soil mulching (M), irrigation regimes (W), growth stage (S)– shown by the three different time periods, and their interactions on mean evapotranspiration (ET), evaporation, and transpiration.

| Treatments | ET (kg/plant) |  |  | Evaporation (kg/pot) |  |  | Transpiration (kg/plant) |  |  |
| --- | --- | --- | --- | --- | --- | --- | --- | --- | --- |
|  | 0–30 days | 31–60 days | 61–98days | 0–30 days | 31–60 days | 61–98 days | 0–30 days | 31–60 days | 61–98 days |
| SHS mulch (N) | 3.66±0.14 | 4.89±0.21 | 5.29±0.22 | 0.19±0.02 | 0.21±0.01 | 0.27±0.01 | 3.48±0.13 | 4.68±0.20 | 5.02±0.22 |
| Bare soil (N) | 3.93±0.14 | 4.59±0.16 | 5.34±0.15 | 0.93±0.11 | 0.93±0.16 | 1.12±0.18 | 3.00±0.21 | 3.66±0.20 | 4.23±0.30 |
| SHS mulch (R) | 1.79±0.14 | 2.07±0.24 | 2.20±0.33 | 0.18±0.40 | 0.20±0.02 | 0.18±0.02 | 1.61±0.13 | 1.86±0.23 | 2.02±0.31 |
| Bare soil (R) | 2.24±0.15 | 2.39±0.17 | 2.40±0.11 | 0.92±0.02 | 0.86±0.08 | 0.68±0.01 | 1.32±0.15 | 1.57±0.22 | 1.74±0.07 |
| **3-way ANOVA** | **ET** |  |  | **Evaporation** |  |  | **Transpiration** |  |  |
| M (df =1) | ns |  |  | 266 (0.0*) |  |  | 25.0 (0.0*) |  |  |
| W (df =1) | 702 (0.0*) |  |  | 5.63 (0.02) |  |  | 500 (0.0*) |  |  |
| S (df = 2) | 33.0 (0.0*) |  |  | ns |  |  | 26.0 (0.0*) |  |  |
| M×W (df = 1) | ns |  |  | ns |  |  | 5.28 (0.03) |  |  |
| M×S (df = 2) | ns |  |  | ns |  |  | ns |  |  |
| W×S (df = 2) | 15.0 (0.0*) |  |  | 3.49 (0.04) |  |  | 7.74 (0.0*) |  |  |
| M×W×S (df = 2) | ns |  |  | ns |  |  | ns |  |  |

Data presented in the upper rows indicate mean values ( ± SE) of total ET, evaporation, and transpiration at three time periods (0–30, 31–60, and 61–98 days after transplanting) for each treatment combination: SHS-mulched mulch under **N** irrigation (SHS mulch-N); bare soil under **N** irrigation (Bare soil-N); SHS-mulched mulch under **R** irrigation (SHS mulch-R); and bare soil under **R** irrigation (Bare soil-R). The lower rows indicate three-way ANOVA statistics for the effects of **M**, **W**, **S**, and their interactions on total ET, evaporation, and transpiration. Significant results are represented by *F* values followed by their respective *p*-values (*p* < 0.05) in parentheses; ns = non-significant results; 0.0\* represents *p*-values < 0.001; df = degrees of freedom.

### Effects of SHS mulching on physiological traits, biomass yield, and fruit yield

The performed two-way ANOVA demonstrated the significant effect of SHS mulching on total evaporation and transpiration (*p* < 0.05) but no significant effect of SHS mulching on the total ET, as presented in **Table 2**. In addition to the changes in ET, evaporation, transpiration, and plant height demonstrated in **Figs. 2–3**, SHS mulching impacted other plant responses such as stomatal conductance and aperture, CCI, TE, fruit yields, HI, fresh mass, dry mass, root xylem vessel, and total root diameter, as presented in **Figs. 4−7**.

**Table 2.** Results of two-way ANOVA test showing effects of soil mulching (M), irrigation regime (W), and their interactions (M × W) on plant traits measured.

| Source of variation | df | Total ET | Total evaporation | Total transpiration | Stomatal conductance | Chlorophyll content index, CCI |
| --- | --- | --- | --- | --- | --- | --- |
| M | 1 | ns | 92.7 (0.0*) | 12.3 (0.0*) | 17.5 (0.0*) | 62.6 (0.0*) |
| W | 1 | 197 (0.0*) | ns | 232 (0.0*) | 23.5 (0.0*) | 17.4 (0.0*) |
| M×W | 1 | ns | ns | ns | ns | ns |
| Error |  | 0.002 | 0.196 | 0.849 | 15810 | 26.584 |

| Source of variation | df | Plant height | Shoot fresh mass | Root fresh mass | Total fresh mass | Shoot dry mass |
| --- | --- | --- | --- | --- | --- | --- |
| M | 1 | 11.7 (0.01) | 51.4 (0.0*) | 15.8 (0.0*) | 53.2 (0.0*) | 21.5 (0.0*) |
| W | 1 | 43.4 (0.0*) | 89.9 (0.0*) | 29.9 (0.0*) | 111 (0.0*) | 80.3 (0.0*) |
| M×W | 1 | ns | 12.5 (0.0*) | ns | 11.2 (0.0*) | 7.65 (0.02) |
| Error |  | 6.468 | 103.310 | 5.971 | 114.8 | 5.619 |

| Source of variation | df | Root dry mass | Total dry mass | Transpiration efficiency, TE | Fruit yields per plant | Harvest Index, HI |
| --- | --- | --- | --- | --- | --- | --- |
| M | 1 | 20.6 (0.0*) | 14.1 (0.0*) | ns | 6.19 (0.03) | 15.6 (0.0*) |
| W | 1 | ns | 36.2 (0.0*) | 45.1 (0.0*) | 23.3 (0.0*) | 12.2 (0.0*) |
| M×W | 1 | ns | ns | ns | ns | ns |
| Error |  | 0.792 | 10.518 | 0.1326 | 1541 | 0.004 |

Data presented are *F* values from two-way ANOVA and *p*-values for significant results ( $p < 0.05$ ) in parentheses; ns = non-significant results. 0.0\* represents $p$ -values $< 0.001$ ; df = degrees of freedom.

**Fig. 4.**
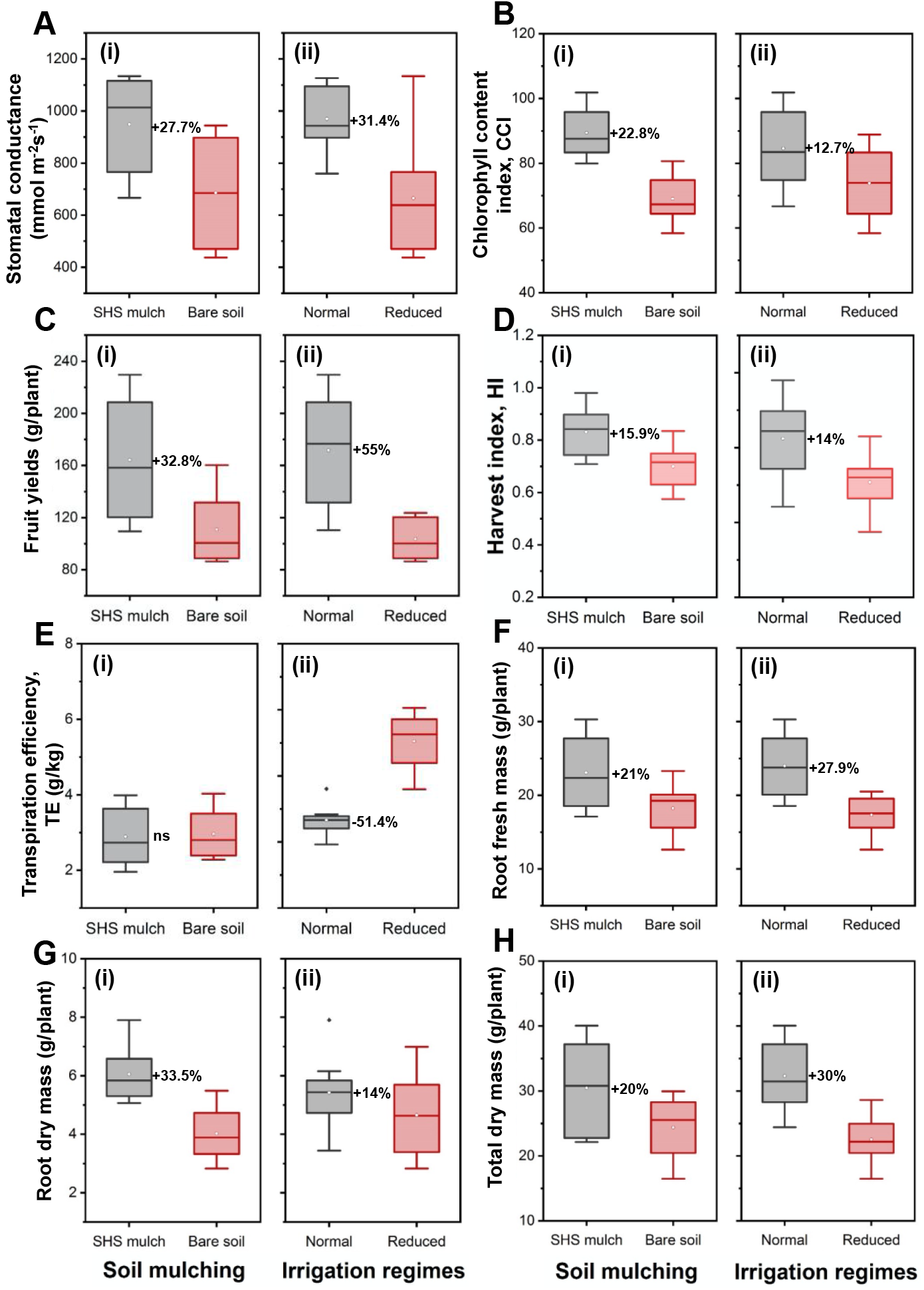
ANOVA results showing effects of (**i**) soil mulching (SHS mulch vs. bare soil) and (**ii**) irrigation regimes (**N** vs. **R**) on (**A**) leaf stomatal conductance; (**B**) CCI; (**C**) fruit yields per plant; (**D**) HI; (**E**) TE; (**F**) root fresh mass; (**G**) root dry mass; and (**H**) total dry mass per plant. Each box represents the data distribution from eight plants (*n* = 8) with the median along the mid-line. The white dot inside the box represents the mean value, the upper and lower sections of the box represent the 25% and 75% confidence intervals, respectively, the whiskers on the box represent the 1.5 interquartile range, and dots outside the box indicate outliers. ns: no significant difference between treatments.

### Main effects of soil mulching and irrigation regimes

A 27.7% higher stomatal conductance was present in SHS-mulched plants than those in bare soil across irrigation regimes (*p* = 0.001; **Fig. 4A-i**). Stomatal conductance was also 31.4% higher under **N** irrigation than under **R** irrigation (*p* < 0.05; **Fig. 4A-ii**). Similarly, leaf CCI was 22.8% higher in SHS-mulched plants than those in bare soil (*p* < 0.05; **Fig. 4B-i**). Irrespective of mulch treatment, leaf CCI was 12.7% higher under **N** irrigation than **R** irrigation (*p* = 0.001; **Fig. 4B-ii**). Fruit yield, defined by the total fruit mass per plant, was 32.8% higher in mulched soil than bare soil (*p* = 0.03; **Fig. 4C-i**) and 55% higher under **N** irrigation than **R** irrigation (*p* < 0.05; **Fig. 4C-ii**).

The relationship between fruit yield and total fresh mass (i.e., the HI) was characterized by 16% higher HI in mulched soil than bare soil (*p* = 0.002; **Fig. 4D-i**). Regardless of the mulch treatment, using **R** irrigation decreased the HI by 14% (*p* = 0.004; **Fig. 4D-ii**). No significant difference in the ratio of shoot dry mass to total water transpired per plant (i.e., the TE) was present between plants grown in mulched and bare soils (*p* = 0.697; **Fig. 4E-i**). However, the TE was 51.4% lower under **N** irrigation than **R** irrigation (*p* < 0.05; **Fig. 4E-ii**).

In terms of biomass allocation, plants in mulched soil had 21% more root fresh mass than those in bare soil (*p* = 0.002; **Fig. 4F-i**); plants under **N** irrigation had 28% more root fresh mass than their **R** irrigation counterparts (*p* < 0.05; **Fig. 4F-ii**). The root dry mass was 33.5% higher in mulched soil than bare soil (*p* < 0.05; **Fig. 4G-i**), while it was 14% higher under **N** irrigation than **R** irrigation (*p* = 0.046; **Fig. 4G-ii**). The total dry mass in mulched soil increased by 20% over that in bare soil (*p* = 0.003; **Fig. 4H-i**), while **N** irrigation resulted in 30% higher total dry mass than under **R** irrigation (*p* < 0.05; **Fig. 4H-ii**).

### Interaction effects of mulching and irrigation regimes

There was also a significant interaction effect of mulching and irrigation on shoot fresh mass, shoot dry mass, and total fresh mass per plant. The shoot fresh mass was 26.6% higher in mulched soil than bare soil under **N** irrigation, whereas under **R** irrigation, the shoot fresh mass was 13% higher in mulched soil than bare soil (**Fig. 5A**; *p* < 0.05, *p* = 0.019, respectively). The shoot dry mass increased in mulched soil by 27% compared with that in bare soil under **N** irrigation; under **R** irrigation, shoot dry mass was 12% higher in mulched soil than bare soil (**Fig. 5B**; *p* = 0.001, *p* = 0.046, respectively). Under **N** irrigation, SHS mulching increased the total fresh mass per plant by 26.5%; under **R** irrigation, the total fresh mass per plant was 13.5% higher in mulched soil than bare soil (**Fig. 5C**; *p* < 0.05, *p* = 0.037, respectively).

**Fig. 5.**
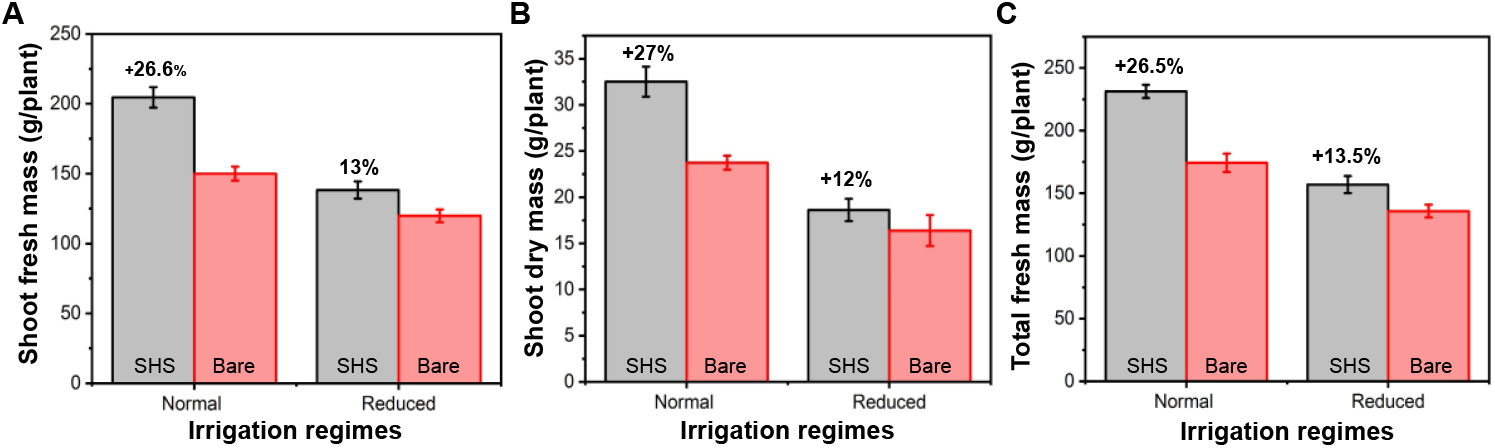
Two-way interaction effects of soil mulching (SHS vs. bare soil) and irrigation regimes (**N** vs. **R**) on (**A**) shoot fresh mass, (**B**) shoot dry mass, and (**C**) total fresh mass per plant. Each bar represents the mean value of four replicates (*n* = 4), and each error bar represents the SE of the means of four samples.

### Leaf stomatal status

Microscopy revealed that the stomatal pores of plants grown in mulched soils (**Fig. 6A-i & iii**) were larger than those in bare soils (**Fig. 6A-ii & iv**); the mean stomatal aperture (area) was 25% larger regardless of the irrigation regime (*p* = 0.0019; **Fig. 6B-i**). Furthermore, the mean stomatal aperture was 34.6% higher under **N** irrigation than **R** irrigation (*p* < 0.05; **Fig. 6B-ii**).

**Fig. 6.**
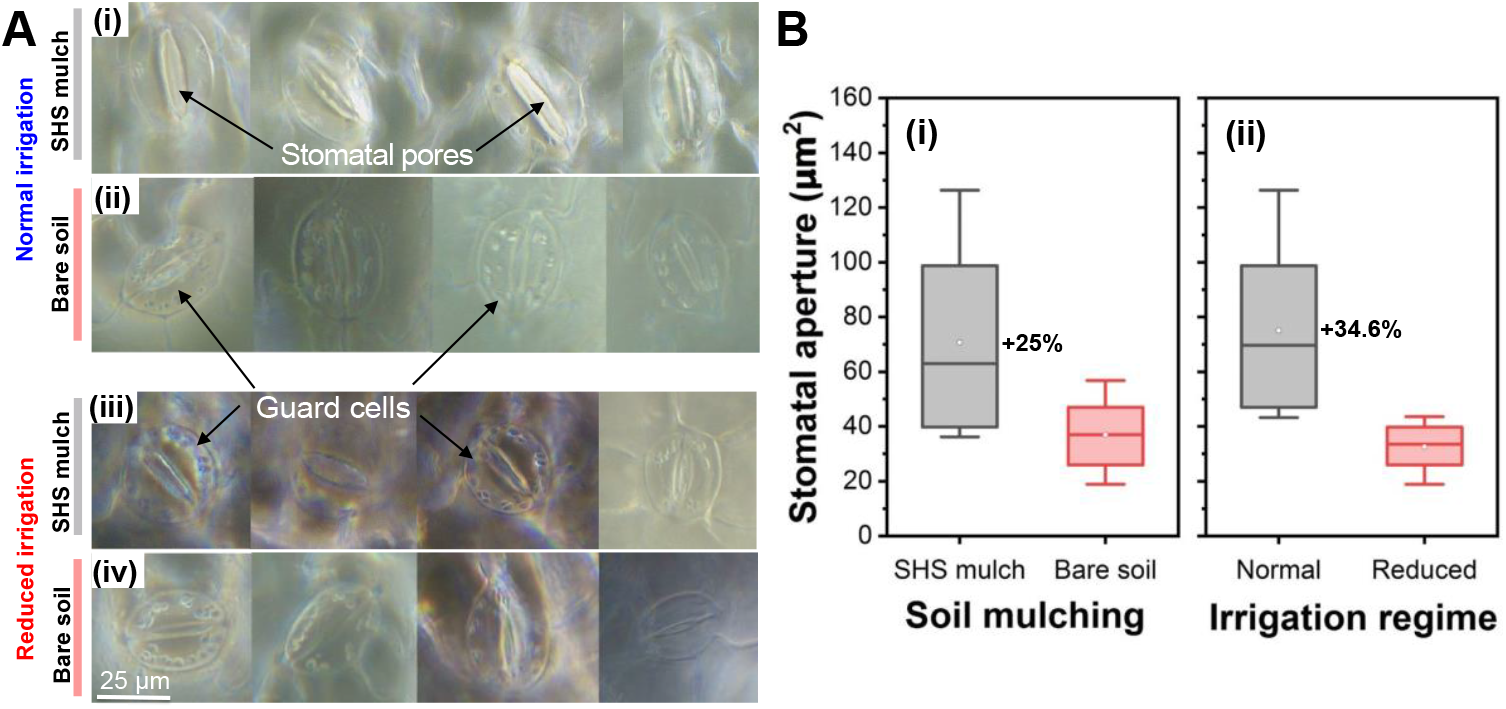
Microscopic analysis of leaf stomatal pores in response to soil mulching and irrigation regimes as on the final date of plant harvest. (**A**) Leica DVM6 microscopic image of leaf stomata for plants grown under **N** and **R** irrigation in (**i, iii**) SHS-mulched and (**ii, iv**) bare soils. (**B**) Boxplots showing the mean stomatal aperture (area) for plants grown in (**i**) SHS and bare soil, and (**ii**) under **N** and **R** irrigation. Each box represents the data distribution from 16 leaf samples (*n* = 16) with the median along the mid-line. The white dot inside the box represents the mean value, the upper and lower sections of the box represent the 25% and 75% confidence intervals, respectively, and the whiskers on the box represent the 1.5 interquartile range.

### Root anatomical structure

Finally, root anatomical features were measured to determine the shoot–root feedback linkages with the observed trends in ET as a function of mulching and irrigation scenarios; these results are summarized in **Figs. 7A-i–7A-iv**. The xylem vessel diameter was 21% larger for roots grown in mulched soil than bare soil (*p* = 0.001; **Fig. 7B-i**). However, the difference in diameter under differing irrigation scenarios was not statistically significant (*p* = 0.343; **Fig. 7B-ii**). Additionally, the root diameter increased in mulched soil by 28% relative to bare soil (*p* = 0.0177; **Fig. 7C-i**), whereas it decreased from **R** irrigation to **N** irrigation by 20% (*p* = 0.0067; **Fig. 7C-ii**).

**Fig. 7.**
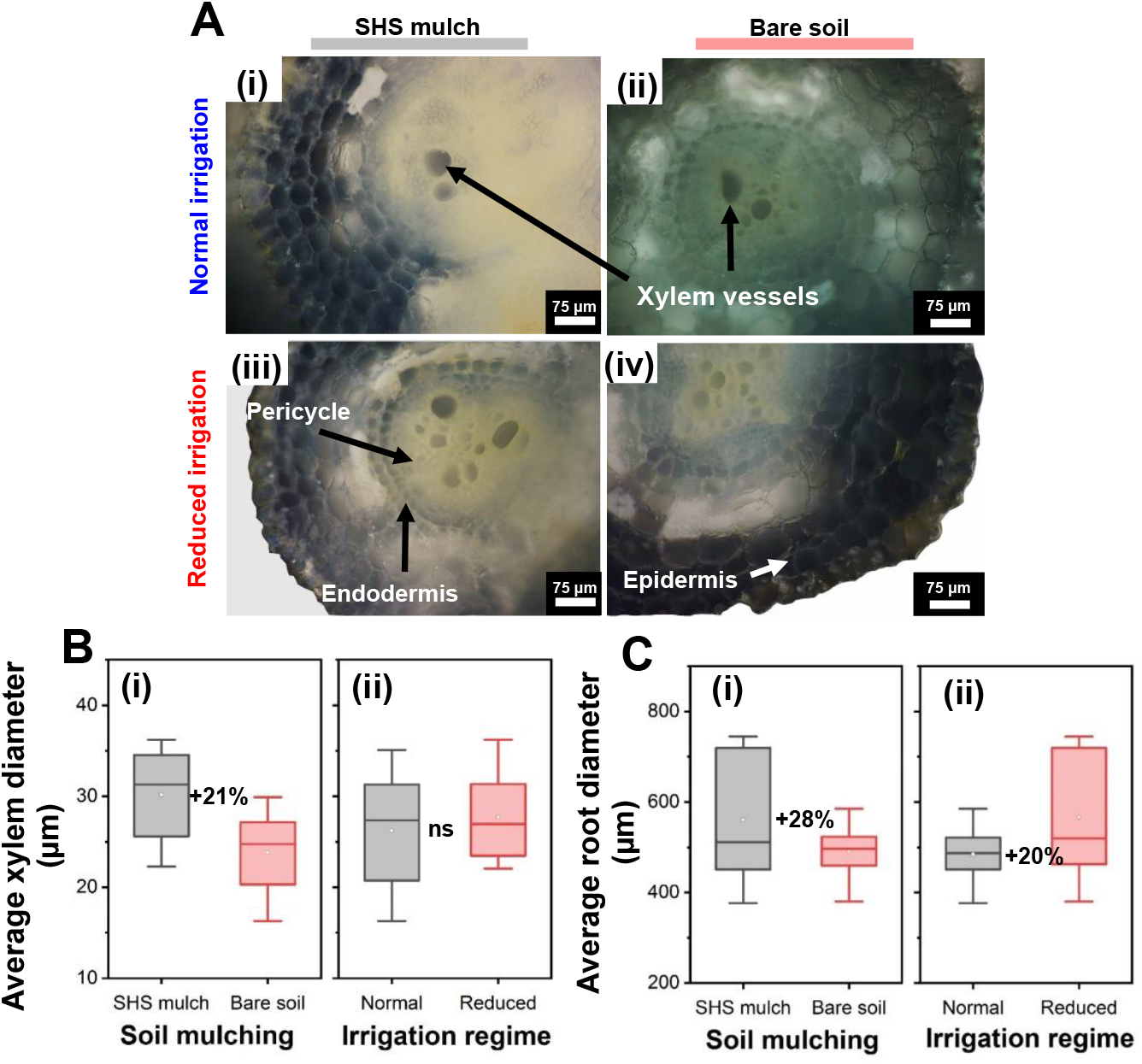
(**A, i–iv**) Cross-sections of tomato roots from contrasting soil mulching and irrigation scenarios. (B-C) Box plots showing the effects of (i) SHS mulching and (ii) irrigation regimes on (**B**) average xylem vessel diameter and (**C**) average root diameter. Each box represents the data distribution from 16 root samples (*n* = 16) with the median along the mid-line. The white dot inside the box represents the mean value, the upper and lower sections of the box represent the 25% and 75% confidence intervals, respectively, the whiskers on the box represent the 1.5 interquartile range, and dots outside the box indicate outliers. ns: no significant difference between treatments.

## DISCUSSION

To clarify the effects of SHS mulching presented on ET fluxes and phenotypic traits of tomato plants grown under N and R irrigation, the mechanistic insights behind SHS mulching and their various impacts on plants and interrelationships are analyzed in this section. The physical mechanism behind the ability of SHS to reduce evaporation from soil is based on the Laplace pressure (i.e., the pressure difference across a curved liquid surface), which depends on the surface tension of water and the curvature of the liquid−air interface (Domingues et al., 2018, Das et al., 2019, Gallo Jr et al., 2021a). When water is supplied through sub-surface irrigation, it rises upward through the soil by capillarity and contacts the SHS layer on the topsoil. The curvature of the air–water interface touching the SHS grains forms a convex shape; the capillary action that drives the rise of water in soils prevents it from wetting the SHS layer, creating a diffusion barrier (Gallo Jr et al., 2021a, Gallo Jr et al., 2021b). By physically covering the topsoil, the mulch also insulates the soil from solar radiation and wind, thereby further reducing the capacity for evaporation losses (Kader et al., 2017, Gallo Jr et al., 2021a). With less evaporation, the soil moisture content rises, allowing for higher plant transpiration.

A significant causal relationship was present between the stomatal conductance and aperture (**Fig. 8A**) and between transpiration and stomatal conductance (**Fig. 8B**), total fresh mass (**Fig. 8C**), total dry mass (**Fig. 8D**), and fruit yield (**Fig. 8E**). These relationships suggest that the soil moisture-retaining function of SHS augments a chain of plant-related responses that affect stomatal opening and closure, which directly influence transpiration rates. Plants can alter their stomatal pore aperture by actively adjusting the turgor pressure within their guard cells (Blatt, 2000), which moderates the exchange of gases such as water vapor, carbon dioxide (CO_2_), and oxygen between the leaf interior and the atmosphere (Kollist et al., 2014, Lawson and Vialet-Chabrand, 2019, Franks and Farquhar, 2007). Enhanced water availability in the soil increases the guard cell turgor pressure, leading to a larger stomatal aperture and increased stomatal conductance of water vapor, which enhances the transpiration and CO_2_ uptake rates for photosynthesis (Bertolino et al., 2019, Condon et al., 2002, Putra et al., 2012). This rationale underpins the observed increase in transpiration (**Figs. 2−3**; **Table 1**), stomatal conductance (**Fig. 4A**), and stomatal aperture (**Fig. 6**) when using SHS mulch.

**Fig. 8.**
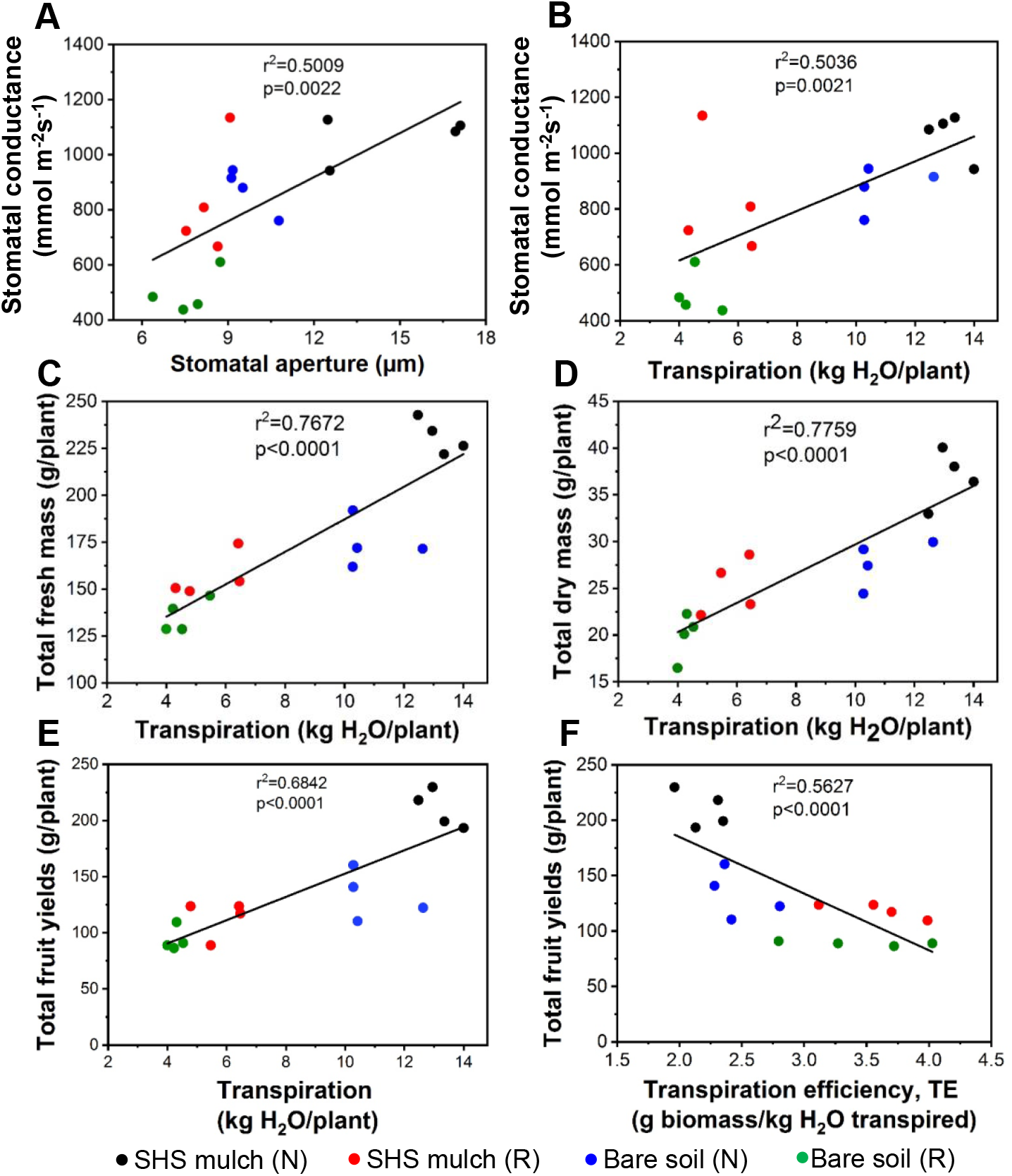
Regression analyses showing the relationship between (**A**) leaf stomatal conductance and stomatal aperture, (**B**) stomatal conductance and total transpiration, (**C**) total fresh mass and total transpiration, (**D**) total dry mass and total transpiration, (**E**) total fruit yield per plant and total transpiration, and (**F**) total fruit yield per plant and TE. Each dot on the graph indicates an individual plant grown in SHS mulch or bare soil under **N** or **R** irrigation.

Aboveground physiological processes, including stomatal responses, are governed by a feedback mechanism linked closely with the hydraulic conductivity of the soil water in the root zone (Putra et al., 2012, Tardieu and Parent, 2017, Zarebanadkouki et al., 2016, Bertolino et al., 2019, Adachi et al., 2010, Chaves et al., 2002). This hydraulic conductivity is dependent on the diameter of the roots and the xylem vessels (Niklas, 1985, Comas et al., 2013). Here, larger xylem vessel and root diameters were present in mulched soil than in bare soil (**Fig. 7**). Consistent with these findings, larger xylem vessel and root diameters have been shown to develop in wheat plants in response to increased SWC caused by mulching (Zhan et al., 2019), increasing WUE and shoot biomass (Kadam et al., 2015). Although small-diameter roots maintain plant growth during a drought, especially at soil depths with available water (Comas et al., 2013), several researchers have suggested that large-diameter roots are more instrumental in resource uptake and plant growth under water scarcity (Larson and Funk, 2016, Kadam et al., 2015). The latter agrees with the observed results; the roots had a larger average diameter under **R** irrigation than **N** irrigation. A larger diameter may promote root penetrative ability or soil exploration in dry soils (Clark et al., 2003) and enhance root longevity (McCormack et al., 2012). These results indicate the important role that root architecture plays in resource acquisition and conservation to facilitate aboveground processes that determine productivity in plants.

The root–shoot hydraulic system conducts nutrients and water via the transpiration stream to the leaves; these nutrients (e.g., nitrogen) support plant growth and the photosynthetic process. Nitrogen availability is related to the total chlorophyll content in plant leaves (Bassi et al., 2018). The higher the chlorophyll content, the more efficient its photosynthetic capacity is, causing increased growth, yields, and biomass (Condon et al., 2004, Haefele et al., 2009). Here, a higher CCI and higher transpiration due to SHS mulching contributed to higher fruit yield and biomass production, leading to a higher HI. The fruit mass per plant was negatively correlated with TE (**Fig. 8F**), indicating that fruit yields increased under low TE. This finding contradicts the proposal that higher TE is critical in increasing crop yields in water-limited environments (Christy et al., 2018, Sinclair, 2012, Rebetzke et al., 2002, Coupel-Ledru et al., 2016, Condon et al., 2004, Vadez and Ratnakumar, 2016). However, TE is a physiologically complex trait that results from a combination of multiple physiological parameters, such as photosynthesis, stomatal conductance, mesophyll conductance (i.e., the diffusion of CO_2_ from sub-stomatal cavities to the sites of carboxylation in the chloroplasts) (Flexas and Medrano, 2002, Flexas et al., 2008), and other conditions that determine carbon balance and growth in plants (Natarajan et al., 2021). Consequently, the conditions responsible for the expression of high TE in one situation may be associated with a low TE in another environment (Sinclair, 2012).

A prolonged stomatal opening and higher stomatal conductance account for lower TE (Flexas et al., 2008); mulched soils retain more SWC and maintain stomatal openings longer than bare soils, leading to a low TE (Balwinder-Singh et al., 2011). However, no significant difference in TE was present between SHS-mulched and bare soils, despite their difference in stomatal conductance. However, lower stomatal conductance was associated with higher TE under **R** irrigation than under **N** irrigation. These findings indicate the trade-off required between stomatal conductance and carboxylation efficiency to improve TE and yields. Unlike increasing the stomatal conductance, increasing the mesophyll conductance increases photosynthesis and TE without increasing transpiration because the CO_2_ diffusion pathway involving mesophyll conductance is not shared with that of transpired water (Flexas et al., 2008, Ouyang et al., 2017, Flexas et al., 2013, Barbour et al., 2010).

Future investigations are needed to examine physiological traits associated with enhanced TE under different environmental conditions. For example, experiments resolving instantaneous leaf gas exchange could help determine photosynthesis and conductance (stomatal vs. mesophyll conductance), whereas measurements of carbon isotope composition (*δ*^13^C; a surrogate trait that integrates TE and variation in root traits and plasticity) could enhance understanding of how aboveground and belowground processes shape plant responses and yield performance under different water regimes. Determining the genetic factors that can maximize transpiration and carbon assimilation to enhance photosynthetic output would complement the water-saving benefit of SHS mulching to improve yields in arid land agriculture.

## CONCLUSIONS

In conclusion, by combining a broad set of techniques to investigate the effects of SHS mulching on ET fluxes and plant responses, this study revealed that adding a thin (1.0-cm thick) layer of SHS dramatically decreased water evaporation from the topsoil, thereby boosting transpiration under both irrigation scenarios studied. The enhanced transpiration, in turn, increased plant biomass and fruit yield. Furthermore, this study facilitated understanding the mechanistic insights behind increased transpiration, biomass, and fruit yield owing to the root–shoot hydraulic processes accounting for enhanced water uptake and stomatal regulation. The quantitative assessment of ET, evaporation, and transpiration presented here will aid crop WUE assessments and field water optimization and management using SHS mulching, irrigation systems design, and agricultural regimes in arid and semi-arid regions.

## ACKNOWLEDGMENTS

The co-authors would like to thank the assistance of the following staff of KAUST Plant Growth Core Labs Facilities: Dr. Richard Soppe, Dr. Muppala, and Mr. John E. Rahmer.

## FUNDING

This research was supported by funding from King Abdullah University of Science and Technology under the award number **BAS/1/1070-01-01**.

## AUTHOR CONTRIBUTIONS

**KO:** Designed and performed the experiments, including soil preparation, planting, harvests, data collection, and analysis; manuscript writing and revision. **AGJ:** Assisted with the experimental design, data analysis, and manuscript revision. **VD:** Collected the evapotranspiration data and assisted in plant harvest and biomass determination. **HM:** Supervised this research and revised the manuscript.

## DATA AVAILABILITY STATEMENT

All data supporting the findings reported in this study are available in the paper. **HM** and **AGJ** have filed a US patent (US20200253138A1) for the SHS used in this work, while **KO** and **VD** have no competing financial interest in this work.

## Notes

### Competing Interest Statement

Two of the co-authors, Himanshu Mishra and Adair Gallo Jr. have filed a US patent (US20200253138A1) for the superhydrophobic sand mulches used in this work, while Kennedy Odokonyero and Vinicius Dos Santos have no competing financial interest in this work.

